# Investigation of white spot syndrome virus (WSSV) infection in wild crustaceans in the Bohai Sea

**DOI:** 10.1101/2020.08.12.247486

**Authors:** Tingting Xu, Xiujuan Shan, Yingxia Li, Tao Yang, Guangliang Teng, Qiang Wu, Chong Wang, Kathy F.J. Tang, Qingli Zhang, Xianshi Jin

## Abstract

The ecological risks of white spot syndrome virus (WSSV), an important aquatic pathogen, has been causing increasing concern recently. A continuous survey on the prevalence of WSSV in the wild crustaceans of the Bohai Sea was conducted in present study. The result of loop-mediated isothermal amplification detection showed that WSSV positivity rates of sampling sites were determined to be 76.73%, 55.0% and 43.75% in 2016, 2017 and 2018, respectively. And the WSSV positivity rates of samples were 17.43%, 12.24% and 7.875% in 2016, 2017 and 2018, respectively. Meanwhile, the investigation revealed that 11 wild species from the sea were identified to be WSSV positive. The WSSV infection in wild crustacean species was confirmed by transmission electron microscopy analysis. The results of this study suggest that WSSV had been colonized in wild species offshore and the impact caused by WSSV to the wild marine ecosystem cannot be ignored.

## 1. Introduction

Up to now, there were nearly 20 viral pathogens that could cause severe epidemics in shrimp (Lightner & Redman, 1998; Thitamadee *et al*., 2016). Among them, the white spot syndrome virus (WSSV) was considered to be the most serious viral pathogen (Flegel & Fegan, 2002), which had caused up to 100% mortality of the farming shrimp in many farms, resulting in significant economic losses (Lightner,1996). The International Bureau of Animal Diseases (OIE) and the Network of Aquaculture Centres in Asia-Pacific (NACA) lists WSSV as one of the aquatic animal viral pathogens that need to be reported.

WSSV is an enveloped, double-stranded DNA virus that belongs to the genus *Whispovirus* of the family *Nimaviridae* (Witteveldt *et al*., 2004). According to previous reports, WSSV had been prevalent in the major shrimp producing countries around the world since it was first discovered in 1992 (Chou *et al*., 1995), including China, Japan, North Korea, Thailand, South Korea, Indonesia, Vietnam, Malaysia, India, Sri Lanka, Bangladesh and the United States (Huang *et al*., 1995; Inouye *et al*.,1994; Momoyama *et al*., 1993; Nakano *et al*.,1993; Takahashi *et al*., 1994; Wang *et al*., 1995; Lightner *et al*., 1999). The prevalence of WSSV caused huge economic losses to the shrimp farming industry of the world. Moreover, in addition to farmed crustaceans, the presence of WSSV in wild shrimps had also been reported by different researchers (Hossain *et al*., 2001; Jang *et al*.,2009; Marques *et al*.,2011; Soo-Jung *et al*., 2010; Gholamhosseini *et al*., 2020; Mondal & Mandal, 2020). However, there have been no systematic investigation on the prevalence of WSSV in marine ecosystems up to now.

In present study, a large-scale investigation on prevalence of WSSV in the wild crustaceans of the Bohai Sea was conducted in 2016 to 2018. Virus detection and confirmation were conducted by using loop-mediated isothermal amplification (LAMP) assay and transmission electron microscopy (TEM).

## 2. Materials and methods

### 2.1 Sample collection

The bottom trawl surveys were carried out at 101 designated sampling sites (Supplementary Table 1) in the Bohai Sea in May and August of 2016-2018. Three to six individuals of the dominant species of marine crustaceans were sampled at every designated sampling site. Whereas, due to the bad sea weather conditions, samples were collected at 59 designed sites finally. Each individual sample was cut along the longitudinal axis, and one part was preserved in 2.5% glutaraldehyde solution (Sinopharm, Beijing, China). The other part was smeared on Whatman FTA Elute cards (GE, Marlborough, MA, USA). The FTA Elute card was dried for 30 min under natural conditions and then sealed and stored at -20 °C for later detection.

**Table 1.** Sampling numbers and WSSV positivity rates of different crustacean species collected in the Bohai Sea in 2016-2018

| **Scientific name** | **2016** | **2017** | **2018** | **WSSV positivity rate** |
| --- | --- | --- | --- | --- |
| *Alpheus distinguendus* | 45 | 23 | 46 | 21.93% (25/114) |
| *Crangon affinis* | 45 | 8 | 40 | 19.35% (18/93) |
| *Palaemon gravieri* | 87 | 16 | 57 | 16.25% (26/160) |
| *Acetes chinensis* | 8 | \ | 36 | 13.64% (6/44) |
| *Leptochela gracilis* | 10 | 2 | 3 | 13.33% (2/15) |
| *Trachypenaeus curvirostris* | 71 | 10 | 72 | 9.74% (15/154) |
| *Alpheus japonicus* | 107 | 28 | 72 | 6.76% (14/207) |
| *Latreutes planirostris* | \ | \ | 4 | 25% (1/4) |
| *Euphausia pacifica* | \ | \ | 2 | 50% (1/2) |
| *Latreutes anoplonyx* | 4 | \ | 1 | 60% (3/5) |
| *Penaeus chinensis* | 2 | 1 | \ | 66.67% (2/3) |
| *Eualus siensis* | \ | \ | 1 | 0 (0/1) |
| *Exopalaemon carinicauda* | \ | 8 | 2 | 0 (0/10) |
| *Penaeus japonicus* | \ | \ | 2 | 0 (0/2) |
| *Metapenaeus joyneri* | \ | \ | 1 | 0 (0/1) |
| *Sicyonia cristata* | \ | \ | 1 | 0 (0/1) |
| *Oratosquilla oratoria* | \ | \ | 2 | 0 (0/2) |
| *Upogebia major* | \ |  | 1 | 0 (0/1) |
| *Lysmata vittata* | \ | 1 | \ | 0 (0/1) |
| **SUM** |  | **820** |  |  |

### 2.2 Detection of WSSV by LAMP

A piece of paper about 3 mm square was cut off from each sampled FTA® cards and was put into in Eppendorf tube of 1.5 mL. The papers carrying the samples were washed following the manufacturer’s protocol. The washed papers were transferred to a microtube filled with 40 μL of TE buffer and then incubated at 95 °C for 5 min for denaturing of nucleic acids captured on the papers. The papers carrying denatured nucleic acids of the samples were used as the template for WSSV LAMP assay. The reaction of WSSV LAMP was performed in a PCR tube, and a microliter of fluorescent dye (GeneFinder™, Bio-V, Xiamen, China) was pre-sealed into the cap of the reaction tube using paraffin. When the amplification of LAMP assay finished, the reaction product was immediately mixed with GeneFinder™ for fluorescence development by inverting the mixture after incubating at 95 °C for 5 min, and the developed green fluorescent color in the reaction tube will be considered as positive.

### 2.3 Transmission electron microscopy (TEM) analysis

For confirmation of WSSV infection in wild crustaceans, WSSV positive samples determined in LAMP assay were chosen for further TEM analysis. The samples preserved in 2.5% glutaraldehyde solution was subjected to further fixation with 1% osmium tetroxide, and dehydrated in a graded ethanol series, then embedded in Spurr’s resin and finally were stained with uranyl acetate and lead citrate following the protocols described previously (Graham and Orenstein, 2007; Zhang *et al*., 2017). Ultrathin sections were prepared on collodion coated grids by the Equipment Center of the Medical College of Qingdao University. All grids were examined in JEOL JEM-1200 electron microscope operating at 80 kV to 100 kV.

## 3. Results

### 3.1 WSSV prevalence in the wild species of the Bohai Sea

A total of 820 samples of wild crustaceans were finally collected from 59 sampling sites in the Bohai Sea during 2016-2018 (Table 1 and Fig. 1). The prevalence of WSSV in samples from each sampling site was investigated by using WSSV LAMP assay firstly. The results of LAMP assay showed that WSSV positivity rates of sampling sites were determined to be 76.73%, 55.00% and 43.75% in 2016, 2017 and 2018, respectively, which indicated that WSSV had distributed in most area of the Bohai sea. The WSSV positivity rates of samples were 17.43%, 12.24% and 7.875% in 2016, 2017 and 2018, respectively. The WSSV positivity rates both in the sampling sites and in the collected samples showed a gradual downward trend from 2016 to 2018 (Fig. 2). Moreover, the results of LAMP assay also showed that the prevalence rate of WSSV in the wild crustacean species of the Bohai Sea was the highest, and the prevalence scope was the widest in 2016 during the investigation of three years. The positive sampling sites were also distributed throughout the Bohai Sea in 2016. In 2017, although the number of sampling sites decreased, the proportion of positive sampling sites was still very high. And the positive sampling sites were distributed in three bays of the Bohai Sea, including Laizhou Bay, Bohai Bay and Liaodong Bay. Interestingly, in 2018, the positive sampling sites were mainly concentrated in Laizhou Bay, while there was only one positive sampling site in Liaodong Bay (Fig. 1).

**Fig.1.**
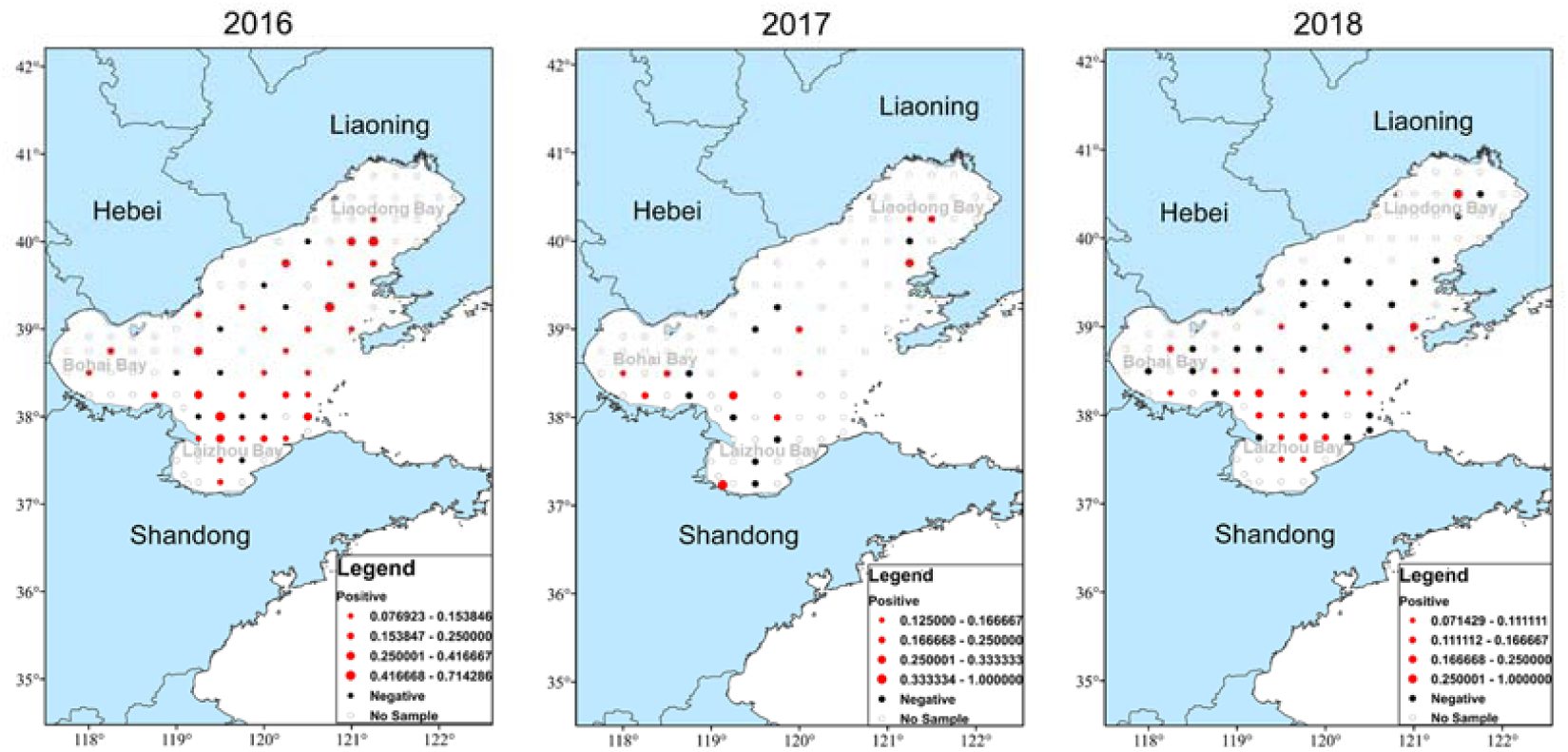
Prevalence rate and prevalence scope of WSSV in the wild crustaceans of the Bohai Sea (2016-2018). The 101 designated sampling sites were showed by solid and hollow spots. The red solid spots indicated that WSSV positive samples were found in the sampling site. The bigger of the spots means the higher of the WSSV prevalence rate in the sampling site. The black solid spots indicated that no WSSV positive samples were found in the sampling site. The hollow spots indicated that no samples were collected in the sampling site.

**Fig. 2.**
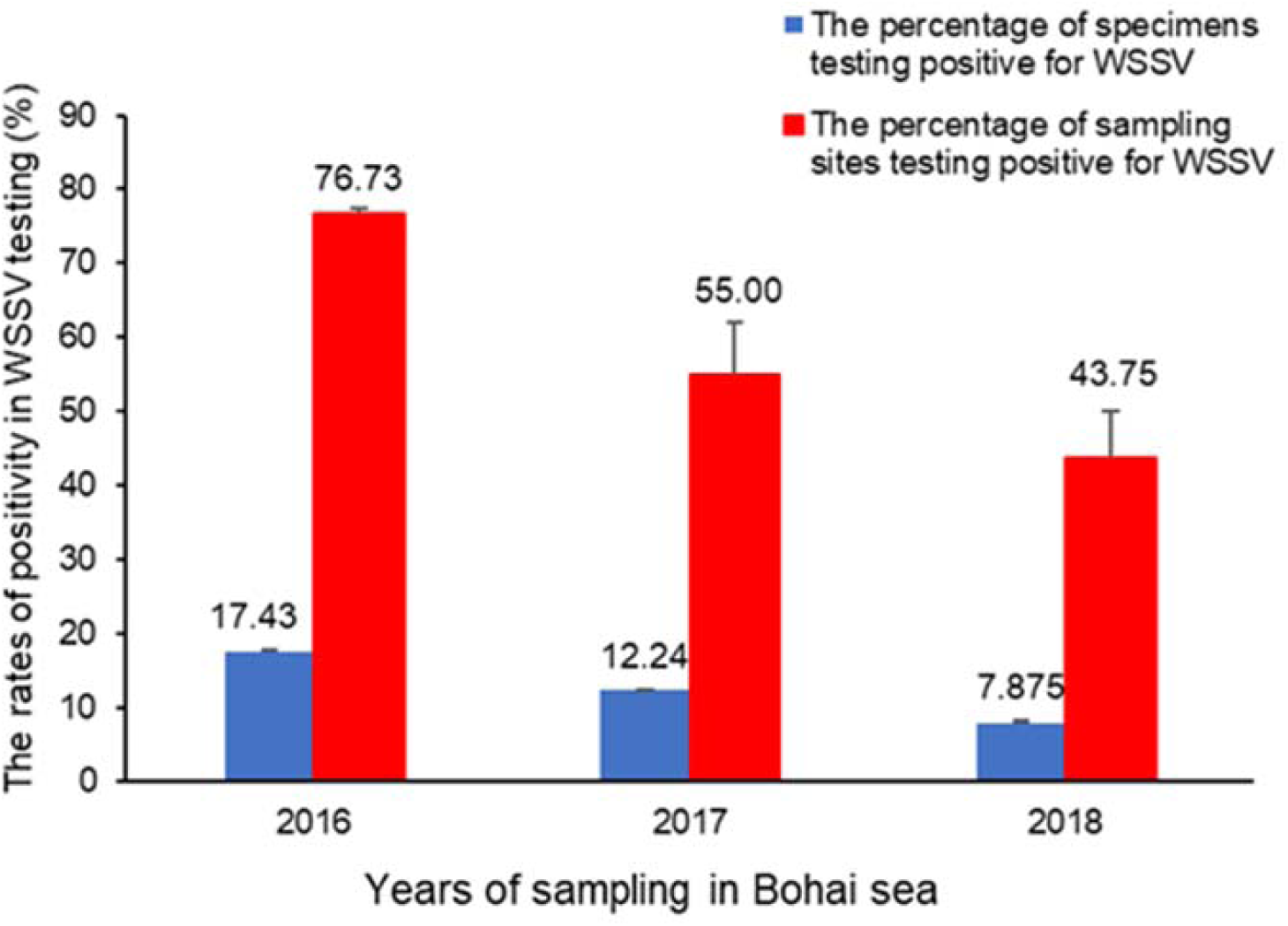
WSSV positivity rates in the sampling sites and in the collected samples in the Bohai Sea (2016-2018).

In 2016-2018, samples of 19 wild crustacean species were collected from the Bohai Sea (Table 1). The results of LAMP detection showed that 11 wild crustacean species, including *Euphausia pacifica, Leptochela gracilis, Latreutes anoplonyx, L. planirostris, Acetes chinensis, Crangon affinis, Palaemon graviera, Alpheus japonicus, A. distinguendus, Trachypenaeus curvirostris, Penaeus chinesis* were determined as positive for WSSV (Table 1).

The WSSV prevalence in the traditional dominant species of crustaceans in the Bohai Sea, including *P. gravieric, A. japonicus, A. distinguendus, A. chinensis, C. affinis, T. curvirostris* and *L. gracilis* were monitored and analyzed in the epidemiological investigations (Fig. 3a). The results showed that the prevalence of WSSV in the dominant species of crustaceans in the Bohai Sea showed a downward trend from 2016 to 2018, except for the *P. graviera* and *T. curvirostris* (Fig. 3b).

**Fig. 3.**
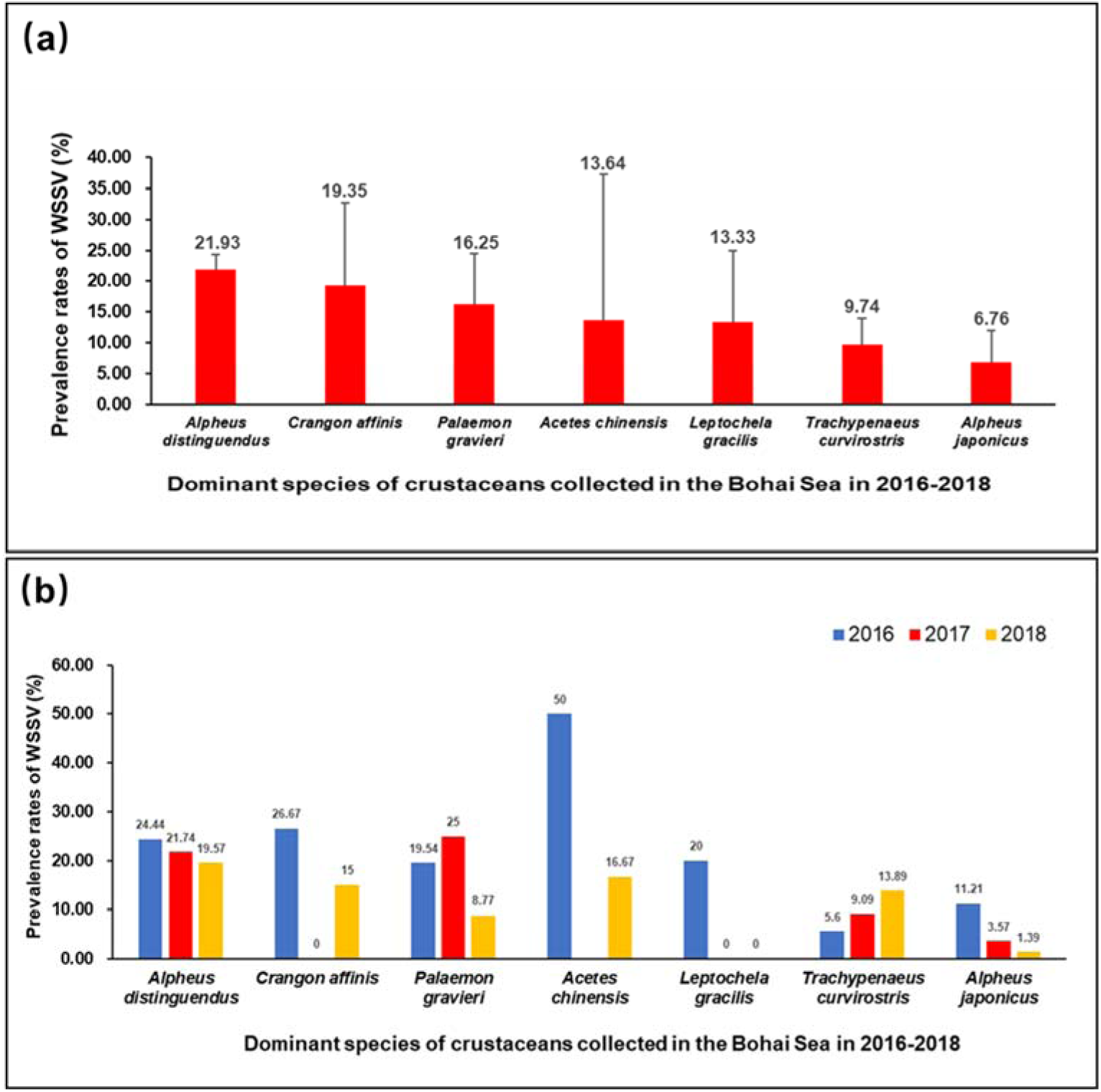
The positivity rate of WSSV in dominant species of crustaceans and the interannual variation of the positivity rate in dominant specie of crustaceans in the survey of the Bohai Sea (2016-2018). (a) The positivity rate of WSSV in dominant species of crustaceans in the survey of the Bohai Sea in 2016-2018. (b) The interannual variation in dominant species of crustaceans in the survey of the Bohai Sea in 2016-2018.

### 3.2 Confirmation of WSSV infection in wild species by TEM

The samples of *T. curvirostris*, one of the dominant species of crustaceans in the Bohai Sea, were chosen for confirmation of WSSV infection in wild species. Under the TEM, a group of enveloped WSSV-like particles with a length of 200 ± 60 nm and a width of 60 ± 15 nm could be observed in the ultrathin sections of muscle of *T. curvirostris* with typical WSSV infection (Fig. 4).

**Fig. 4.**
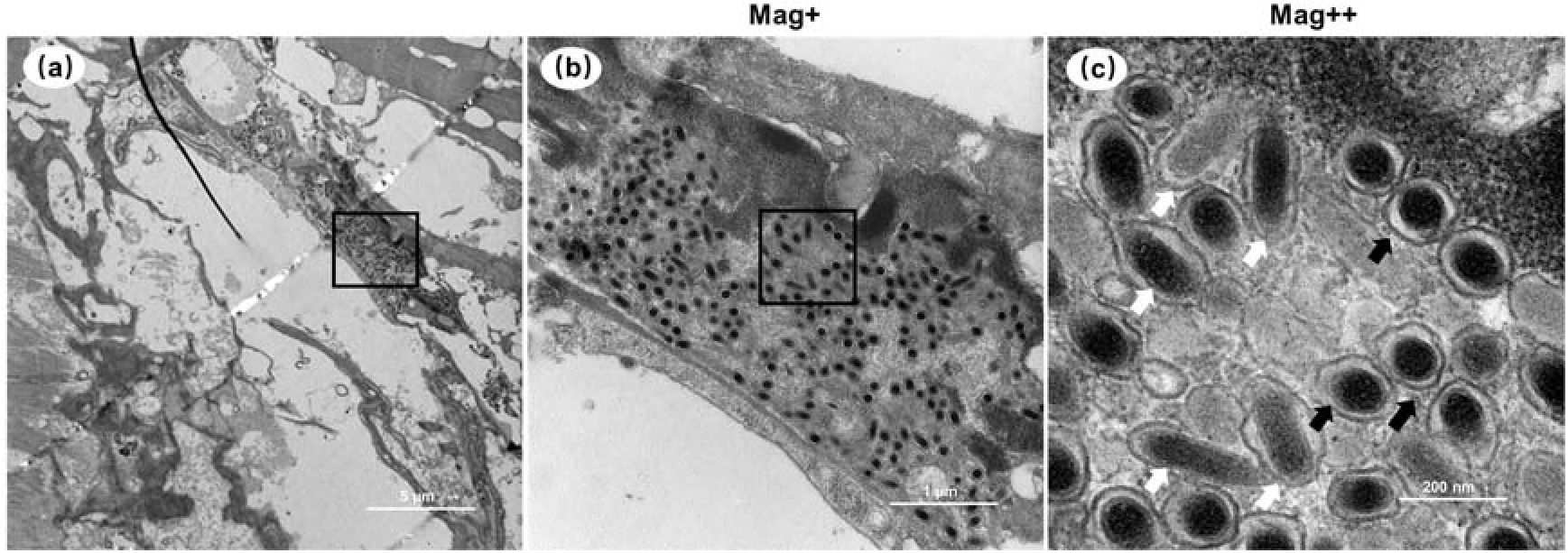
Transmission electron micrographs of WSSV virions in sub-epidermal epithelial cells of wild *Trachypenaeus curvirostris*. (a) TEM of the sub-epidermal epithelial cells of *T. curvirostris*. (b) Magnified micrograph of the partial zone in the black frame in (a). Note that the scattering distribution of WSSV-like particles can be observed. (c) Magnified micrograph of the partial zone in the black frame in (b). black arrows: cross-sections of virions; white arrows: longitudinal sections of virions. Scale bars = (a) 5 µm, (b) 1µm, and (c) 200 nm.

## 4. Discussion

WSSV, a member of the *Nimaviridae*, had spread to most shrimp farming countries and regions around the world since it was reported in 1992, and its ongoing pandemic has caused significant economic losses to the global shrimp farming industry (Vlak *et al*., 2004; Chou *et al*., 1995; Huang *et al*., 1994; OIE, 2003). In China, WSSV prevention was more successful recently because of implementation of strict quarantine policy for the origin of seedlings and the widely using of WSSV-free post larvae of shrimp by the farmers. In addition, the control of WSSV transmission in ponds level was also highly effective to a great extent by exclusion of potential viral carriers from the ponds by prohibiting the use of live bait. Nowadays, researchers have gradually turned their attention to the study of the impact of WSSV on wildlife. In this study, we conducted a continuous survey on the prevalence of WSSV in the wild species of the Bohai Sea from 2016 to 2018.

The results of investigation on prevalence of WSSV in the Bohai Sea showed that WSSV positivity rates of sampling sites and samples were high and even reached 76.73% and 17.43% in 2016, respectively, which indicated that WSSV had been widely prevalent in almost the entire Bohai Sea. It was report that WSSV might originated from certain wild species in natural waters (Rozenberg *et al*., 2015), and the dispersal of WSSV from infected shrimp farms to the marine environment might also occurred in some areas (Mijangos-Alquisires *et al*., 2006). Bohai Sea, surrounded by the Shandong Peninsula and the Liaodong Peninsula, is China’s inland sea. Aquaculture activities around the Bohai Sea was active in the past decades, so the seawater and biological exchanges between coastal ponds and sea areas were very frequent. The high WSSV positivity rates of samples and sampling sites in Bohai sea might be due to the pathogens exchange occurring with material exchange between the coastal terrestrial ponds and offshore waters. Another possibility of high WSSV positivity rates in the Bohai Sea might cause by the large quantity of stock enhancement (or the artificial proliferation) of the crustaceans around the Bohai Sea in the past years (Cui *et al*., 2002; Wang, 2020). The result of investigation indicated that both the WSSV positive sites and WSSV prevalence showed a downward trend year by year in the Bohai Sea during 2016 to 2018, which could be attributed to the decreasing of spillover and dispersal of WSSV from shrimp farmed ponds, because the local government executed the strict quarantine policy of seedlings and local farmers were guided to use WSSV-free post larvae of shrimp. This result seemed corroborating previously reported speculation that improving the surrounding environment along the seashore appeared to be the most effective way to reduce the negative impact of aquaculture pathogens in the ocean (Groner *et al*., 2016; Zhu *et al*., 2019).

The results of investigations on the WSSV positivity rates of samples in different wildlife species from the Bohai Sea in 2016-2018 showed that 11 wild species collected in the Bohai Sea were identified as WSSV positive by LAMP assay. The presence of WSSV virions in the sub-epidermal epithelial cells of wild *T. curvirostris* was further confirmed by TEM analysis, indicating that CMNV infection did happen in the wild crustacean species in the Bohai Sea. Meanwhile, high prevalence of WSSV had been found in samples of dominant crustacean species in the Bohai Sea. The dominant crustacean was the major prey for a variety of predators for a long period (Dou *et al*., 1992; 1993; Zhang *et al*., 2012; We *et al*., 2018) and played key role in maintaining the ecological equilibrium of the Bohai Sea (Deng *et al*., 1988; Liu *et al.*, 2000; Zeng *et al.*, 2017; Wu *et al*., 2012). So, it could be deduced that WSSV prevalence in the major dominant crustacean species might threat the ecological balance and the crustacean stock enhancement of the Bohai Sea in certain degree.

## 5. Conclusions

In summary, infection and prevalence of WSSV in major dominant crustacean species were proved in the surveyed coastal water based on the systematic investigation of wild crustaceans in the Bohai Sea. The results demonstrated that WSSV had been colonized in wild species offshore and the impact caused by WSSV to the wild marine ecosystem cannot be ignored.

## Acknowledgements

The authors would like to thank Mr. Fangqun Dai and the staffs in the research vessel for them generous help in sampling. This work was supported by the National R&D Program of China (2017YFC1404503), Projects of International Exchange and Cooperation in Agriculture, Ministry of Agriculture and Rural Affairs (MARA) of China-Science, Technology and Innovation Cooperation in Aquaculture with Tropical Countries along the Belt and Road, Project of Species Conservation from the MARA-Marine fisheries resources collection and preservation, and Central Public-interest Scientific Institution Basal Research Fund, YSFRI, CAFS (NO. 20603022019003; 20603022020005).

## Author Contributions

Qingli Zhang and Xianshi Jin designed the experiments. Xianshi Jin and Xiujuan Shan design and funded the bottom trawl surveys. Tingting Xu executed the surveillance. Tingting Xu, Qingli Zhang, and Xiujuan Shan analyzed data. Tao Yang, Guangliang Teng, and Qiang Wu help to collect the samples in the survey. Tingting Xu, Yingxia Li, and Chong Wang conducted the molecular assays of the samples. Qingli Zhang conducted the TEM analysis. Tinging Xu, Qingli Zhang, and Xiujuan Shan wrote the manuscript. All authors interpreted the data, critically revised the manuscript for important intellectual contents and approved the final version.

