## Supplementary table for "Investigation of white spot syndrome virus (WSSV) infection in wild crustaceans in the Bohai Sea"

### Supplementary Table 1

Supplementary Table 1. List of the sampling sites in the Bohai Sea

| Sampling site | Longitude | Latitude | Sampling site | Longitude | Latitude | Sampling site | Longitude | Latitude |
|---|---|---|---|---|---|---|---|---|
| **162** | 121 | 40.75 | **2294** | 119 | 39 | **4762** | 118 | 38.25 |
| **184** | 120.833 | 40.5 | **2294** | 119 | 39 | **4851** | 118.25 | 38.25 |
| **194** | 121 | 40.5 | **2353** | 119.25 | 39.167 | **4862** | 118.5 | 38.25 |
| **251** | 121.25 | 40.75 | **2394** | 119.5 | 39 | **4951** | 118.75 | 38.25 |
| **262** | 121.5 | 40.75 | **2451** | 119.75 | 39.25 | **4962** | 119 | 38.25 |
| **283** | 121.25 | 40.5 | **2494** | 120 | 39 | **5051** | 119.25 | 38.25 |
| **294** | 121.5 | 40.5 | **2551** | 120.25 | 39.25 | **5083** | 119.25 | 38 |
| **351** | 121.75 | 40.75 | **2594** | 120.5 | 39 | **5094** | 119.5 | 38 |
| **383** | 121.75 | 40.5 | **2651** | 120.75 | 39.25 | **5151** | 119.75 | 38.25 |
| **394** | 122 | 40.5 | **2694** | 121 | 39 | **5183** | 119.75 | 38 |
| **474** | 122.167 | 40.5 | **2752** | 121.25 | 39.25 | **5194** | 120 | 38 |
| **574** | 120.167 | 40 | **3432** | 118 | 38.917 | **5251** | 120.25 | 38.25 |
| **594** | 120.5 | 40 | **3451** | 117.75 | 38.75 | **5262** | 120.5 | 38.25 |
| **641** | 120.583 | 40.25 | **3462** | 118 | 38.75 | **A** | 119.08 | 38.1 |
| **652** | 120.833 | 40.25 | **3483** | 117.75 | 38.5 | **5283** | 120.25 | 38 |
| **662** | 121 | 40.25 | **3494** | 118 | 38.5 | **5294** | 120.5 | 38 |
| **683** | 120.75 | 40 | **3521** | 118.25 | 38.917 | **6094** | 119 | 37.5 |
| **694** | 121 | 40 | **3532** | 118.5 | 38.917 | **6151** | 119.25 | 37.75 |
| **751** | 121.25 | 40.25 | **3551** | 118.25 | 38.75 | **6162** | 119.5 | 37.75 |
| **762** | 121.5 | 40.25 | **3562** | 118.5 | 38.75 | **6183** | 119.25 | 37.5 |
| **783** | 121.25 | 40 | **3583** | 118.25 | 38.5 | **6194** | 119.5 | 37.5 |
| **794** | 121.5 | 40 | **3594** | 118.5 | 38.5 | **6251** | 119.75 | 37.75 |
| **851** | 121.75 | 40.25 | **3621** | 118.75 | 38.917 | **6262** | 120 | 37.75 |
| **862** | 122 | 40.25 | **3651** | 118.75 | 38.75 | **6283** | 119.75 | 37.5 |
| **883** | 121.75 | 40 | **3662** | 119 | 38.75 | **6294** | 120 | 37.5 |
| **1094** | 119.5 | 39.5 | **3683** | 118.75 | 38.5 | **6334** | 120.5 | 37.8333 |
| **1151** | 119.75 | 39.75 | **3694** | 119 | 38.5 | **6351** | 120.25 | 37.75 |
| **1183** | 119.75 | 39.5 | **3751** | 119.25 | 38.75 | **6372** | 120.1667 | 37.5833 |
| **1194** | 120 | 39.5 | **3794** | 119.5 | 38.5 | **7213** | 119.083 | 37.3333 |
| **1251** | 120.25 | 39.75 | **3851** | 119.75 | 38.75 | **7251** | 119.25 | 37.25 |
| **1294** | 120.5 | 39.5 | **3894** | 120 | 38.5 | **7262** | 119.5 | 37.25 |
| **1351** | 120.75 | 39.75 | **3951** | 120.25 | 38.75 | **7351** | 119.75 | 37.25 |
| **1394** | 121 | 39.5 | **3994** | 120.5 | 38.5 | **B** | 119.13005 | 37.23356 |
| **1451** | 121.25 | 39.75 | **4051** | 120.75 | 38.75 | **/** | / | / |
